# Humans anticipate the consequences of motor control demands when making perceptual decisions between actions

**DOI:** 10.1101/2024.11.14.623637

**Authors:** Élise Leroy, Éric Koun, David Thura

## Abstract

Animals, including humans, are often faced with situations where they must decide between potential actions to perform based on various sources of information, including movement parameters that incur time and energy costs. Consistent with this fact, many behavioral studies indicate that decisions and actions show a high level of integration during goal-directed behavior. In particular, motor costs very often bias the choice process of human and non-human subjects facing successive decisions between actions. However, it appears as well that depending on the design in which the experiment occurs, the effect of motor costs on decisions can vary or even vanish. This suggests a contextual dependence of the influence of motor costs on decision-making. Moreover, it is not currently known whether or not the impact of motor costs on perceptual decisions depend on the difficulty of the decision. We addressed these two important issues by studying the behavior of healthy human subjects engaged in a new perceptual decision-making paradigm in which the constraint level associated with the movement executed to report a choice was volitionally chosen by the participants, and in which the difficulty of the perceptual decision to make continuously evolved depending on their motor performance. The results indicate that the level of constraint associated with a movement executed to express a perceptual decision strongly impacts the duration of these decisions, with a shortening of decisions when these are expressed by demanding movements. This influence appears most important when the decisions are difficult, but it is also present for easy decisions. We interpret this strategy as an adaptive way to optimize the participants’ overall rate of success at the session level.

## INTRODUCTION

Although choices are always ultimately expressed via actions, decision-making and motor control are most often studied separately from each other (Gold and Shadlen, 2007; Shadmehr and Krakauer, 2008; Franklin and Wolpert, 2011; Lee et al., 2012). Recent behavioral studies, including ours, indicate however that decisions and actions are closely linked, sharing economic principles and showing a high level of integration during goal-directed behavior (Shadmehr, 2010; Gallivan et al., 2018; Carland et al., 2019; Shadmehr et al., 2019; Wispinski et al., 2020; Gordon et al., 2021).

For instance, it has been proposed that movement selection, preparation, and execution are parameterized following economical rules, varying depending on utility estimation: high valued options lead to faster reaction times and movement speed, and high-perceived effort discounts an option value, leading to slower reaction and longer movements (Shadmehr et al., 2010; Haith et al., 2012; Choi et al., 2014; Morel et al., 2017). If the sensory information guiding the choice is weak and the decision takes time, humans and monkeys shorten the duration of the movements expressing this choice (Thura et al., 2014; Thura, 2020; Herz et al., 2022). If the context in which the task occurs encourages fast and risky decisions, humans and monkeys report these choices with faster movements compared to when the task is performed in a slow speed-accuracy context (Thura et al., 2014; Thura, 2020; Herz et al., 2022; Carsten et al., 2023; Saleri and Thura, 2024).

Conversely, several studies have demonstrated that motor costs influence decision-making as well, whether choices only rely on movement properties (Cos et al., 2011; Morel et al., 2017; Michalski et al., 2020; Canaveral et al., 2024), options value (Pierrieau et al., 2021; Grießbach et al., 2022) or perceptual stimuli (Burk et al., 2014; Lepora and Pezzulo, 2015; Marcos et al., 2015; Hagura et al., 2017). In our lab, we have demonstrated that humans and monkeys decide faster and/or with less precision in order to focus on their actions when the movement expressing a choice is demanding (Reynaud et al., 2020; Saleri Lunazzi et al., 2023) or time consuming (Saleri Lunazzi et al., 2021; Saleri and Thura, 2024).

Although these studies indicate a significant level of integration between perceptual decision-making and motor control processes, with notably a significant influence of motor costs on perceptual decision-making, two questions need to be addressed. First, it is unknown whether the impact of the motor costs on decision-making depends on the obligation for the subjects to express choices in a difficult motor condition structured in a succession of many consecutive trials. Indeed, in the studies cited above (Reynaud et al., 2020; Saleri Lunazzi et al., 2021, 2023; Saleri and Thura, 2024), the demanding motor condition was always imposed on the subjects, in dedicated blocks of trials. However, it is possible that if the subject deliberately chooses to express a choice in a demanding motor condition, on a trial by trial basis, to obtain a larger reward for example, the impact of these motor demands on the decision process differs. In support of such a possibility, a recent result indicates for instance that the level of movement effort may not influence perceptual decisions when that effort is explicitly felt and integrated by human subjects (Manzone and Welsh, 2023), suggesting that the results of the experiments mentioned above may be specific to how motor costs are manipulated. Secondly, it is currently unknown to what extent the impact of the motor control demands on decision-making depends on the difficulty of the choice to make. It is for instance possible that subjects sacrifice their decisions in favor of more demanding motor control only when the decision is hard but that when the decision is easy, the influence of the motor context is less pronounced or even disappears.

We investigated these two questions by studying the behavior of healthy human subjects engaged in a new perceptual decision-making paradigm in which the constraint level of the movement executed to report a choice was chosen by the participants, and in which the difficulty of the perceptual decision to make continuously evolved depending on their motor performance. This design therefore allowed us to test the effect of motor constraints on perceptual decision-making when these constraints were volitionally chosen by the subjects, and it offered at the same time the possibility of testing this effect on multiple levels of decisional difficulty manipulated in a gradual manner.

## RESULTS

Thirty-two healthy human participants performed a new behavioral paradigm (Fig. 1) during a single experimental session. The goal of the subjects was to accumulate a total of 200 points to complete the session. To earn points, they had to choose at the beginning of each trial the constraint level of the movement executed to report a perceptual decision: either a demanding arm movement, in terms of motor control, potentially earning 5 points if accurately executed, or an easy movement, earning only 1 point if accurately executed. After making that choice, they had to make the corresponding perceptual decision and report it by executing the arm movement toward a visual target. Crucially and unknown to the subjects, the coherence of the visual stimulus was linearly and inversely indexed to the number of accumulated points during the session, progressively increasing the difficulty of each perceptual decision. The points (5 or 1) that the subjects chose to engage at the beginning of each trial were lost in case of a movement error, i.e. if they failed to reach the chosen target and stay in it within the required time windows, but not in case of a wrong perceptual decision.

**Figure 1.**
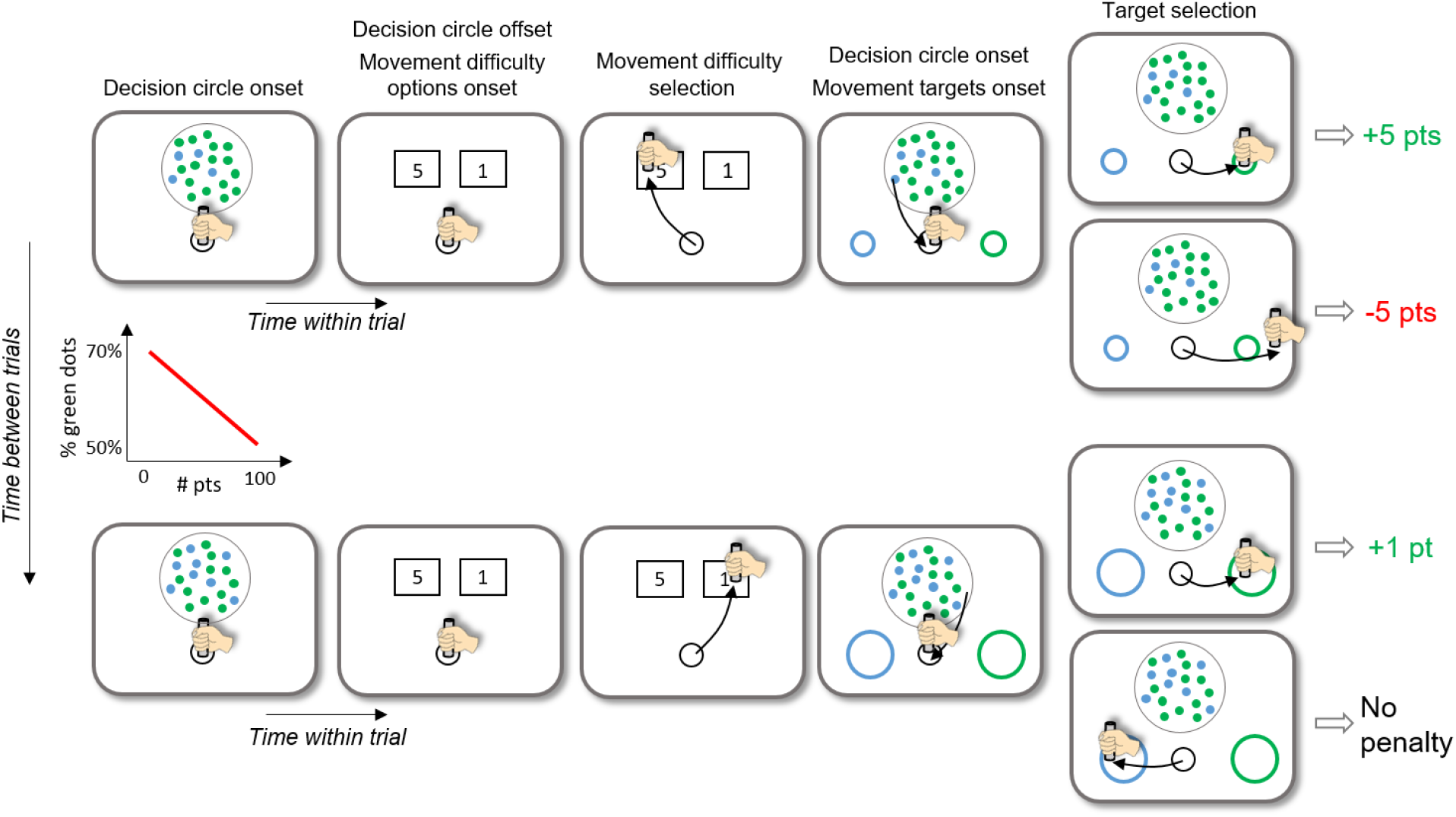
The top row illustrates the time course of a trial at the beginning of the session. Each trial starts when the subject brings with her/his dominant hand a handle and maintains it still in a start circle. The decision circle containing 100 blue and green tokens is first displayed for 300ms to inform the subject about the difficulty of the perceptual decision to make later in the trial. At this stage the dominant color among the tokens (i.e. the coherence) is not informative of the correct target to select at the end of the trial. The proportion of tokens of the dominant color is 77% on the first trial of the session. The decision circle disappears and the movement constraint options are then displayed. In this example the subject chooses “5” (by moving the handle in the rectangle surrounding the number 5), which corresponds to a demanding movement to execute, i.e. toward a small visual target (ø = 1.25cm). Once selected, the subject leaves the motor constraint option rectangle and comes back to the start circle. The decision circle containing the 100 tokens, and the blue and green movement targets then appear. The dominant color among the tokens (with the same proportion as at the beginning of the trial) now determines the correct target to select. The subject reports the decision by moving the handle in the target whose color corresponds to her/his choice. The subject earns the amount of points she/he chose (“5” in this example) if she/he accurately reaches to the correct target. She/he loses the points if she/he executes an inaccurate movement, regardless of the chosen side. After the first trial, the coherence within the decision circle evolves from trial to trial, being linearly and inversely indexed to the number of points accumulated during the session. As a consequence, at the end of the session (bottom row), when the subject gets close to 200 points, the coherence in the visual stimulus is low (proportion of tokens of the dominant color equal to 51%) and the decision difficulty is high. As illustrated in this example, we assumed that in this situation, subjects would choose an easy movement (the “1” rectangle), executed toward a large target (ø = 3.75cm) more frequently than at the beginning of the session, when the decisional effort was low and the number of points to accumulate to complete the session was high. Regardless of the movement constraint level chosen by the subject, if she/he selects the wrong target (decision error) with an accurate movement, points are not deducted.

This design allowed us to assess the impact of the motor control required to express a perceptual decision on participants’ decision-making performance, for decisions whose difficulty varied in small steps from an easy level to a difficult level, and when the demanding motor control condition was deliberately chosen by the subject from one trial to the next to potentially gain more points.

### General observations

Among subjects who performed the task (n=32), the median number of trials to reach 200 points across the population was 184, with a large variability between subjects (min = 87; max = 426; SD = 84 trials). The median proportion of demanding movement choices during a session was 62%, again with high variability between subjects (min: 5%; max: 100%; SD = 34%). With this design we expected to observe a mixture of choices of the constraint level of the movements to be performed to express the perceptual decisions, regardless of the coherence of the decision stimulus. This is because the demanding movements would be chosen to gain more points, perhaps more at the beginning of the session when the number of points to accumulate to complete the session is high and the perceptual decisions are the easiest, while the easy movements would be chosen to guarantee the obtaining of points in case of correct perceptual decisions. However, we observed that out of 32 participants, 9 almost did not vary their movement difficulty choices through the session (1/32 subject chose the easy option in more than 95% of the trials, 8/32 chose that option in less than 5% of trials). In the following analyses, we excluded those 9 subjects who systematically chose the same level of motor constraint level through their experimental session, as they were likely either insensitive (for the 8 subjects who chose the difficult option in more than 95% of the trials) or too sensitive (for the subject who chose the easy option in more than 95% of the trials) to the decisional and/or motor difficulties manipulated in the experiment.

### Effect of motor control demands (i.e. target size) on subjects’ motor behavior

We first verified that the two constraint levels of motor control required to report perceptual decisions did indeed impact the motor behavior of the remaining 23 subjects. To do this, we analyzed the precision and duration of their reaching movements as a function of these two levels of constraint. As expected, we found that participants’ movement accuracy was lower when they reported their perceptual decisions by moving toward the small targets compared to when they had to make a movement toward a large target (median accuracies at the population level: 60% versus 97%, Chi-square test for independence on the population: χ^2^ = 1004, p < 0.0001; Chi-square tests for independence on individual subjects, 21/23 with p < 0.05, Fig. 2, left panel). We also observed that the majority of subjects’ movements (whether accurate or not) were slower, in terms of duration, when executed toward a small target than toward a large target (Wilcoxon rank-sum tests on individual subjects, 14/23 with p < 0.05, Fig. 2, right panel), even if the difference of movement duration between the two motor conditions is not significant at the population level (median durations: 630ms vs. 563ms, Chi-square test for independence on the population: χ^2^ = 1.5, p = 0.13). Given these results, we make the assumption in the following paragraphs that movements executed toward the small targets were more demanding, in terms of motor control, compared to movements executed toward the large targets.

**Figure 2.**
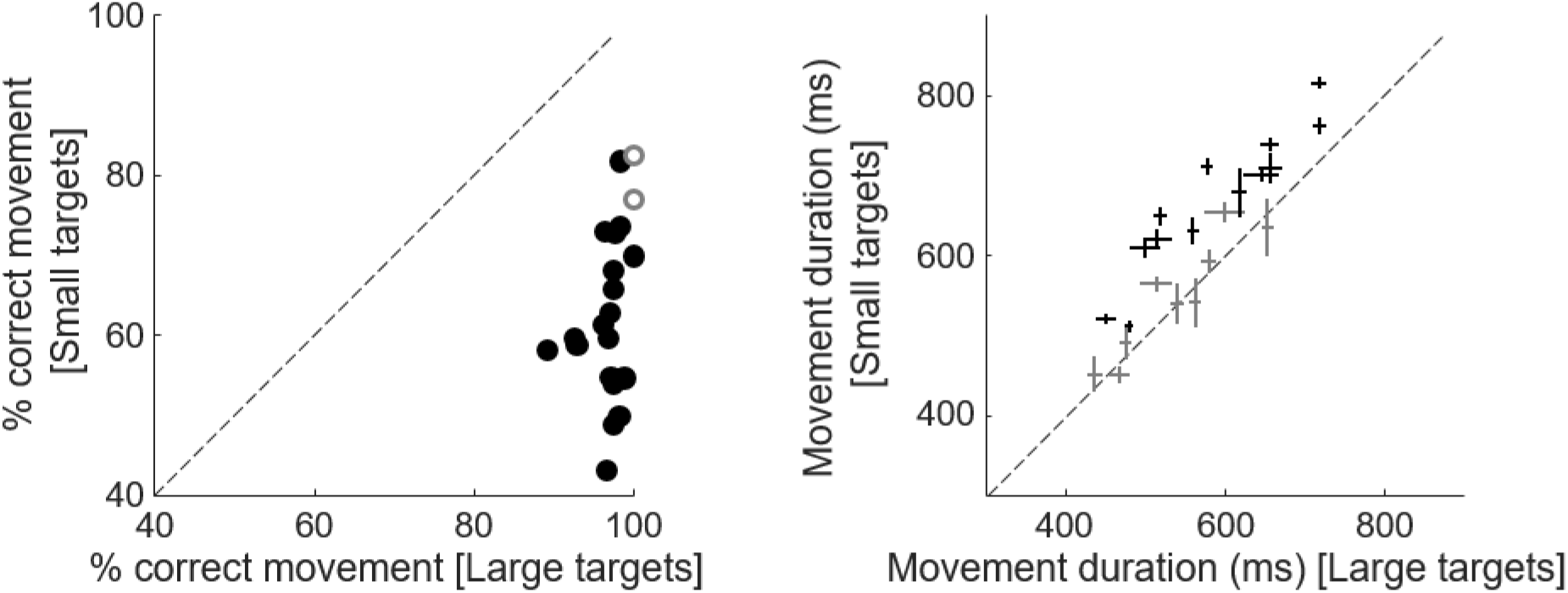
*Left panel*: Comparison of subjects’ movement accuracy as a function of movement constraint level (difficult movement/small targets: ordinate; easy movement/large targets: abscissa). Circles illustrate individual subjects’ data. Filled black circles highlight subjects for which the difference between conditions is statistically significant (Chi-squared test, p<0.05). *Right panel*: Comparison of subjects’ movement duration (whether accurate or not) as a function of movement constraint level (difficult movement/small targets: ordinate; easy movement/large targets: abscissa). Crosses illustrate individual subjects’ medians ± SE. Black crosses highlight subjects for which the difference between conditions is statistically significant (rank-sum test, p<0.05).

### Effect of motor control demands on subjects’ decision behavior

We then analyzed the impact of this demanding level of motor control on the perceptual decisions of the 23 participants. To do this, we first analyzed the accuracy and duration of their perceptual decisions as a function of the motor condition in which decisions were made, regardless of the difficulty of these decisions. We found at the population level that participants’ perceptual decision accuracy was similar whether they were reported with reaching movements executed toward small or large targets (medians: 93.4% versus 90.4%, Chi-square test for independence on the population: χ^2^ = 1.9, p = 0.16). Only two subjects were significantly more accurate to decide when choices were expressed with difficult movements (Chi-square tests for independence on individual subjects, 2/23 with p < 0.05, Fig. 3, left panel). By contrast, we observed that the level of motor control demand strongly impacted the duration of the perceptual decisions preceding the execution of movements executed to report these choices. Indeed, participants were overall faster to decide (accurately or not) when the subsequent movements were demanding compared to when they were easier (median durations: 657ms vs. 739ms, Chi-square test for independence on the population: χ^2^ = −3.4, p < 0.0001). This effect was robust at the individual level, as the effect was significant for the vast majority of subjects (Wilcoxon rank-sum tests on individual subjects, 17/23 with p < 0.05, Fig. 3, right panel).

**Figure 3.**
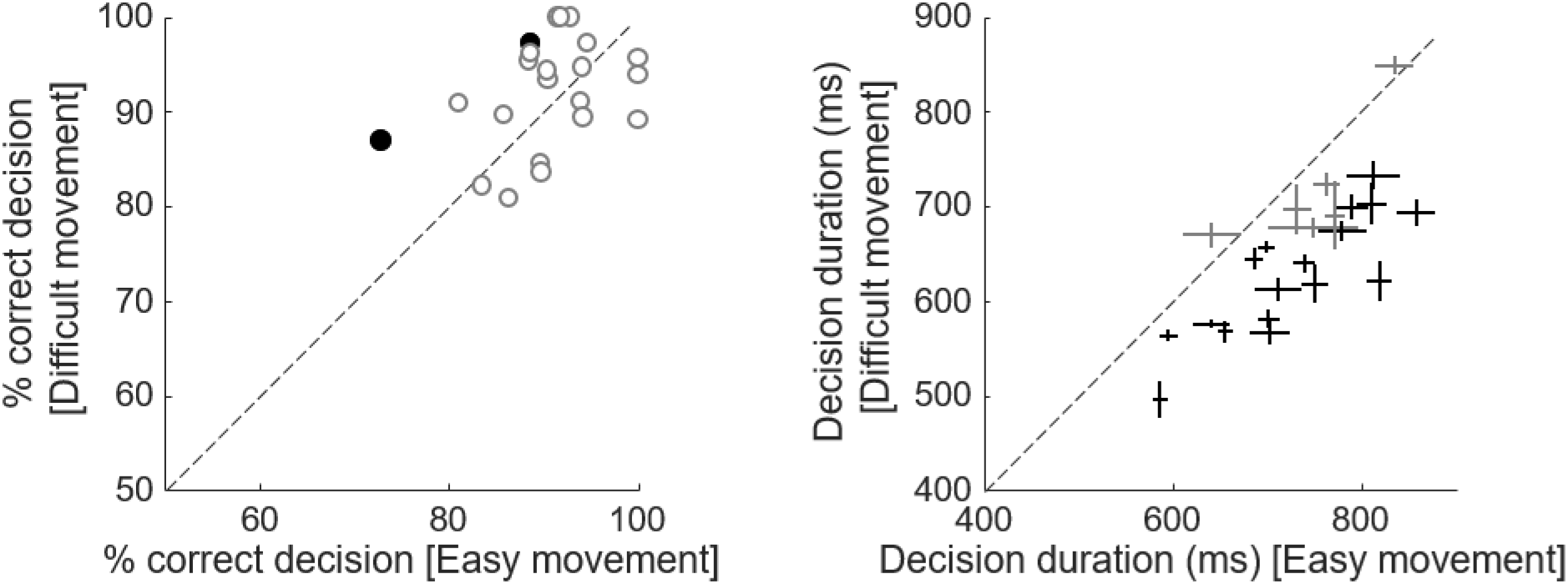
*Left panel*: Comparison of subjects’ perceptual decision accuracy as a function of movement difficulty (difficult movement/small targets: ordinate; easy movement/large targets: abscissa). Same convention as in Fig. 2, left panel. *Right panel*: Comparison of subjects’ perceptual decision durations (correct or not) as a function of movement difficulty (difficult movement/small targets: ordinate; easy movement/large targets: abscissa). Same conventions as in Fig. 2, right panel.

The above results indicate that the level of motor control required to express perceptual decisions, deliberately chosen by individuals, impacts the duration of these decisions preceding the movements. To answer the second question addressed in the present study, namely whether this impact of motor costs on decision-making depends on the difficulty of decisions, we analyzed the behavior of the subjects according to the evolution of their decisional and motor performances in the task.

Our goal with this experimental design was to obtain trials in which subjects would volitionally choose to report their perceptual decisions with easy or difficult movements, while gradually increasing the difficulty of these perceptual decisions in small increments. Because the coherence in the decision stimulus continuously varied from trial to trial, we normalized the number of trials performed by each subject by chronologically grouping them in 10 quantiles. As shown in figure 4A, the first 10% of trials were trials for which the coherence of the decisional stimulus was the highest (because subjects’ scores were the lowest) and thus perceptual decisions were the easiest; Conversely, the last 10% of trials were the trials for which the coherence of the decision stimulus was the lowest, and thus decisions were the most difficult.

**Figure 4.**
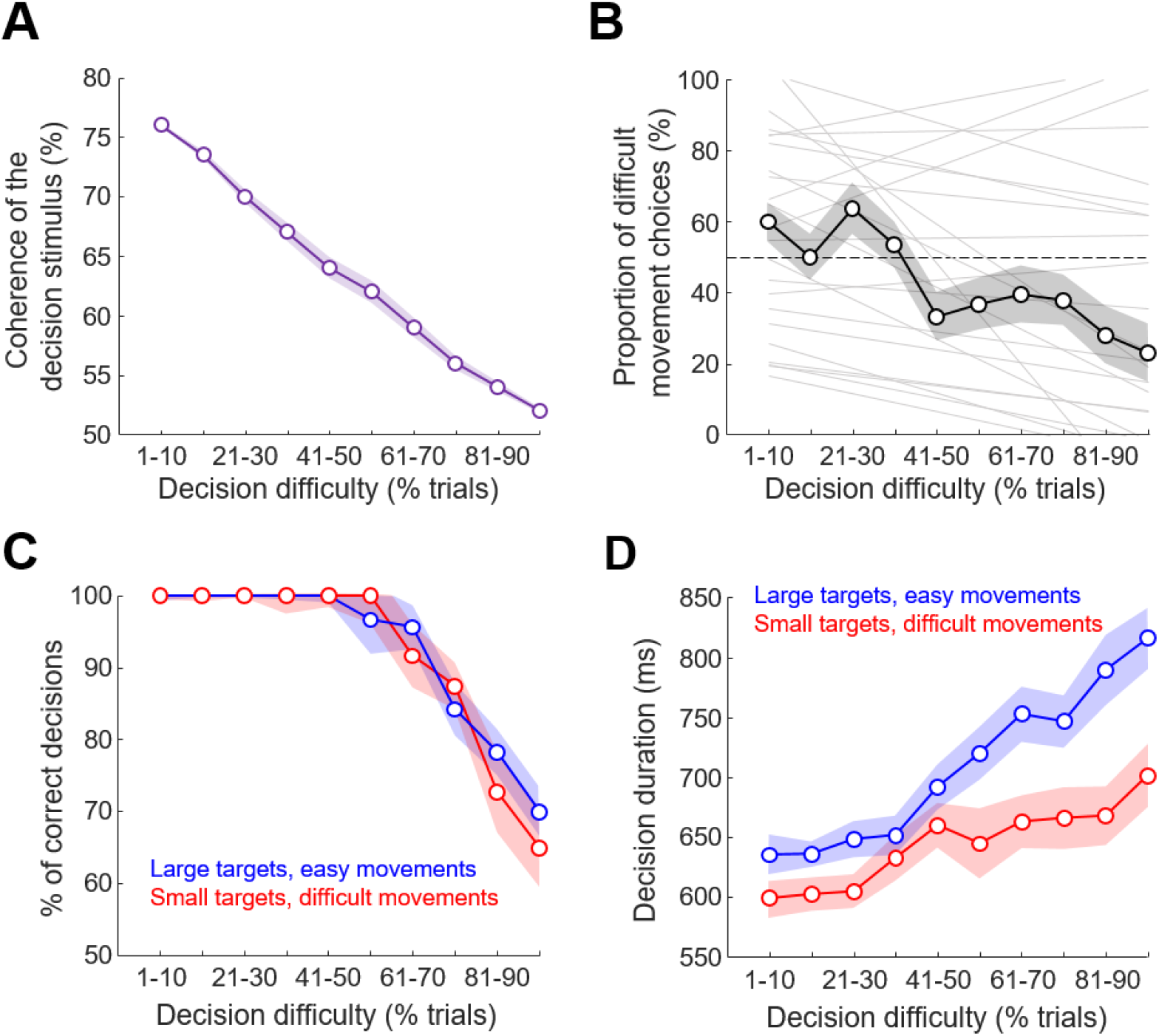
*A*. Relationship between the number of completed trials (averaged ± SE, shaded area, across the population) and the coherence of the decision stimulus. Coherence is defined as the ratio between the number of tokens of each color in the decision stimulus. It is expressed as the percentage of tokens of one color compared to the other. Trials are sorted chronologically and a normalization is performed by grouping them in 10 quantiles. *B*. Proportion of demanding movement choices as a function of decision difficulty. As in *A*, trials are sorted chronologically and normalized by grouping them in 10 quantiles. Because the coherence of the decision stimulus strongly co-varies with the number of completed trials, trial number is a proxy of the decision difficulty. Open circles illustrate the median values (± SE, gray shaded area) at the population level, and light gray lines illustrate linear regressions through the data for each individual subject. *C*. Median (± SE, colored shaded area) proportion of correct perceptual decisions across subjects, as a function of decision difficulty, with trials sorted as a function of the motor condition chosen by the subjects (blue: large targets/easy movement; red: small targets/difficult movement). *D*: Perceptual decision durations as a function of decision difficulty, with trials sorted as a function of the motor condition chosen by the subjects (blue: large targets/easy movement; red: small targets/difficult movement). Same convention as in C.

We found at the population level that the proportion of difficult movement choices did not significantly vary depending on the level of decision difficulty (Kruskal-Wallis test, χ^2^ = 11.2, p = 0.26), despite the fact that a tendency for a decrease of that proportion with the increase of decision difficulty is visible (Fig. 4B). Indeed, at the individual level, we found that the evolution of the perceptual decision difficulty influenced the proportion of movement difficulty choices in 17 out of 23 subjects (Chi-squared tests for independence, p < 0.05). Among them, the vast majority (13/17) overall decreased their proportion of “small targets” choices with the increase of the perceptual decision difficulty (Fig. 4B).

We then analyzed the decision-making behavior of the participants as a function of the difficulty of the perceptual decision and as a function of the level of motor control required to report these choices by performing analyses of covariance (ANCOVAs). As expected, the proportion of correct decisions across the population significantly decreased depending on the number of trials performed during the session (i.e. as a function of the decreasing coherence in the stimulus, see Fig. 4A), regardless of the motor control condition (ANCOVA, Trials: F=199, p < 0.0001; Movement difficulty: F=2.5, p = 0.12; Movement difficulty × Trials: F=0.78, p=0.38, Fig. 4C). As expected too, we observed that the duration of decisions (whether correct or not) increased as a function of the number of trials performed during the session (Trials: F=32.3, p < 0.0001, Fig. 4D). As mentioned above (Fig. 3, right panel), we also found a strong effect of the motor condition on the duration of the perceptual decisions, with decisions being longer when reported via movements executed toward large targets (Movement difficulty: F=104, p < 0.0001, Fig. 4D). But interestingly, we found that this effect was not dependent on the difficulty level of the perceptual decisions, even if a trend for a more pronounced impact when decisions are difficult is observed (Movement difficulty × Trials: F=3.79, p = 0.052, Fig. 4D).

## DISCUSSION

The present study was designed to test the impact of motor demands on the accuracy and duration of perceptual decisions whose difficulty was manipulated in a gradual manner, and when costly movements were volitionally chosen by human subjects from one trial to the next to gain more points. Our study indicates that the level of constraint associated with a movement executed to express a perceptual decision strongly impacts the duration of these decisions. This influence appears most important when the decisions are difficult, but it is also present for easy decisions. In the following paragraphs, we discuss these results in the context of recent work on decision and action interactions, and we propose an interpretation of these data in terms of behavior optimization.

### Motor constraints influence perceptual decision-making

Recent behavioral studies indicate that decisions and actions are closely linked, showing a high level of integration during goal-directed behavior (for reviews, see Shadmehr, 2010; Gallivan et al., 2018; Carland et al., 2019; Shadmehr et al., 2019; Wispinski et al., 2020; Gordon et al., 2021). On the one hand, the properties of movement, such as speed and duration, are influenced by decision-making characteristics (Thura et al., 2014; Thura, 2020; Herz et al., 2022, Carsten et al., 2023; Saleri and Thura, 2024), and the selection, preparation, and execution of movements are parameterized following the same economical rules as those which govern decisional processes (Shadmehr et al., 2010; Haith et al., 2012; Choi et al., 2014; Morel et al., 2017).

On the other hand, numerous behavioral studies have now provided strong support for action-based (or embodied) models of decision-making that hypothesize that action alternatives and their associated properties, including costs, are incorporated into the choice process (Cisek, 2007; Lepora and Pezzulo, 2015). These behavioral studies tested the impact of motor constraints on decision-making using different designs. Very often, a specific motor cost, itself manipulated in different ways, is associated with each option proposed to participants during each trial, and the basis for choosing varies depending on the paradigm. For example, decisions may be based primarily on the motor properties of movements, with one option being costlier than the other in terms of biomechanical cost or energy expenditure on each trial (Cos et al., 2011, 2014; Morel et al., 2017), sometimes during ongoing actions (Michalski et al., 2020; Grießbach et al., 2022; Canaveral et al., 2024). In other cases, the movements performed to express a choice carry different costs but the decision is primarily based on perceptual cues (Burk et al., 2014; Marcos et al., 2015; Hagura et al., 2017) or rewards (Pierrieau et al., 2021; Grießbach et al., 2022) associated with each of the two possible options. In most of these scenarios, the manipulation of motor costs influences the choice of target selected by participants before or during movement execution.

There is, however, evidence that questions a systematic effect of motor costs on decision making, suggesting a task and/or parameter specific aspect to this phenomenon. For example, in a recent study by Manzone and Welsh, the authors demonstrate that when motor costs are clearly and explicitly felt by participants (which is not necessarily the case in other similar studies (Marcos et al., 2015; Hagura et al., 2017)), this explicit effort has a reduced or even absent impact on perceptual decision making (Manzone and Welsh, 2023). In the study of Pierrieau and colleagues, the authors report that when the rewarded target carries the highest motor cost, reaching movements executed within 350ms of reaction time are biased toward the other target, but this effect disappears if participants takes longer to react (Pierrieau et al., 2021). Finally, when comparing the effects of motor costs between a manual movement task and a walking task within the same participants, Grießberg and colleagues observed that the motor costs bias on perceptual decisions only weakly transfer between tasks (Grießbach et al., 2023). Together, these results point to a contextual influence of action effort and costs on perceptual decision-making.

In our own recent experiments, we manipulated the motor condition in which perceptual decisions were made by human or non-human subjects. Specifically, the two options offered to the participants to report their choices (based on changing visual evidence) carried broadly the same motor cost, but this cost varied in dedicated blocks of trials imposed on the subjects. In this case, whether for human subjects or for monkeys, we observed a strong impact of the motor condition on the precision and duration of the perceptual decisions taken before the arm movement was executed to express these choices (Reynaud et al., 2020; Saleri Lunazzi et al., 2021; Saleri and Thura, 2024).

Given the design-related contrasted results of the experiments mentioned above when each movement option carries a specific cost, we could wonder whether in our experimental condition where the two motor targets are associated with the same motor costs, the change of context, notably the voluntary choice of the subjects to execute a more or less costly movement to gain more or fewer points, and this possibly from one trial to another and not in blocks of several dozen trials, could have consequences on the consideration of these motor costs during the perceptual decision-making process. In support of action-based decision models though, we observed that not only do motor costs influence decision-making when motor constraints are voluntarily chosen by the subjects, often from one trial to another and not in blocks, but that this influence persists whatever the difficulty of the decision, suggesting a very close and robust link between motor parameters expressing a perceptual decision and the perceptual decision process.

### Participants seek to preserve their rate of reward

The prospect of executing a more or less demanding movement therefore strongly and systematically modulates the duration of decisions in the present experiment. But how can we explain the meaning of this modulation, namely a shortening of decisions when these are expressed by constrained movements? One might have expected to observe the opposite, i.e. slower and more precise decisions when these choices are expressed by costly movements. Indeed, an influent theory proposed that when an action is costly and requires effort, the overall motivation to behave is reduced, leading to not only slower movements, but also longer reaction times (Mazzoni et al., 2007; Shadmehr et al., 2019). Another interpretation of an identical result would be to consider that the participants sought to optimize the efforts invested in a costly behavior. Indeed, faced with a demanding movement, it is conceivable that they wanted to ensure that they made a good decision by extending the duration of the deliberation, longer deliberation being usually associated with better performance in stable coherence decision paradigms.

However, this is not what we observed. None of the 23 subjects had significantly longer decision times when the movements performed to express a choice were constrained (see Fig. 3, right panel). Our interpretation of the direction of this modulation is that participants sought to optimize the total duration of their response, namely the duration of the decision added to that of the movement. Indeed, in the experiment described in this report, the duration of each trial is entirely dependent on the subject’s behavior, who therefore has control over this duration and over her/his speed-accuracy trade-off strategy. By choosing constrained movements to potentially gain more points, participants possibly integrated that these movements requiring a higher degree of motor control would be slower, in terms of duration, than less constrained movements (Fig. 2, right panel, and see Fig. S1 for the analysis of movement parameters, including duration, between the two motor conditions as a function of decision difficulty). Consequently, subjects possibly sought to compensate for this additional time devoted to movement by shortening the duration of their perceptual decisions, regardless of the difficulty of these decisions. This allows them to maximize their success rate, that is, the number of successful trials, minus the effort associated with those trials, divided by the time required to perform those trials. As several studies have shown, success rate is a parameter that human subjects and animals seek to optimize, more than performance per se, when faced with a succession of decisions and actions (Shadlen et al., 2008; Bogacz et al., 2010; Balci et al., 2011; Carland et al., 2019; Shadmehr et al., 2019). Figure S2 illustrates the response time (decision and movement) of the subjects as a function of the difficulty of the decisions and as a function of the motor condition. Although the response time increases with the increase in the difficulty of the decisions, there is no significant effect of the motor condition on this overall response time. This result is consistent with a mechanism of compensation of motor time costs by the duration of the decisions. Interestingly, this shortening of the decisions had no impact on the precision of these decisions (Fig. 4C), which reinforces the idea that this strategy was beneficial and adaptative in terms of optimizing the success rate. This observation is also compatible with the idea that decisions are primarily based on information from a relatively short time window (Uchida et al., 2006; Yang et al., 2008; Chittka et al., 2009). For simple color discrimination, this can be as short as 30ms (Stanford et al., 2010), but even in much more difficult tasks, it appears to be on the order of 100-300ms (Kiani et al., 2008; Price and Born, 2010).

This supposedly preponderant role of the cost of time on the behavior of the subjects does not mean that effort played no role. Indeed, as mentioned above, the success rate integrates the notion of effort and energy expenditure in its equation. In a recent study, we showed that just like time resources, energy resources can be transferred between decision-making and motor processes, by prioritizing the most critical process for the behavior in question (Leroy et al., 2023). But it also appears that effort does not influence decision-making as robustly as time in an experiment specifically designed to dissociate the impact of both cost types on humans decisional strategy (Saleri Lunazzi et al., 2021). This is possibly because unlike time (Myerson and Green, 1995; Shadmehr et al., 2010), effort, considered in its broad definition, is not always considered as a cost (i.e. the effort paradox, Inzlicht et al., 2018). Even if many experiments of voluntary reaching have shown that when given a choice, humans tend to prefer actions that carry the least biomechanical costs over the more effortful ones (Cos et al., 2011; Marcos et al., 2015; Hagura et al., 2017; Pierrieau et al., 2021; Canaveral et al., 2024), other studies have nevertheless shown that energy optimization is not systematically sought by participants (Kistemaker et al., 2010; Morel et al., 2017; Moskowitz et al., 2023). So, considering that in these particular scenarios where subjects are faced with a multitude of decisions between successive (arm or eye) movements, and that the time and the number of correct trials can be optimized to complete the session as quickly as possible, a notion of success rate that seems to strongly influence behavior, it is time more than effort that most systematically influences the strategy of the participants.

### Conclusions

Taken together, the results of the present study and our past studies indicate that human subjects are able to anticipate the consequence of a demanding movement to execute, when this supplementary cost has been deliberately chosen on a trial-by-trial basis to potentially earn more points and complete the session faster, by shortening the duration of the perceptual decisions preceding the execution of the arm movements. This consideration of motor costs in decision-making does not depend on the difficulty of the choices, and does not impact their precision. We interpret this time compensation strategy as an adaptive way to optimize their overall rate of success at the session level.

## METHODS

### Participants

Thirty-two healthy human subjects (median age ± SD: 25.5 ± 4.2; 26 self-identified as females, 6 as males; 25 right handed) participated in this study. All subjects gave their written informed consent before starting the experiment. The ethics committee of Inserm (IRB00003888, IORG0003254, FWA00005831) approved the protocol on June 7th 2022. All methods were performed in accordance with the relevant guidelines and regulations. Each participant was asked to perform one experimental session. They received a monetary compensation (10 euros per completed session) for participating in this study. All participants also performed another version of the task described in the present report, on a different day, for a study designed to test the hypothesis that the management of effort-related energy resources is shared between decision-making and motor control (Leroy et al., 2023).

### Experimental apparatus

The subjects sat in a comfortable armchair and made planar reaching movements using a handle held in their dominant hand. A digitizing tablet (GTCO CalComp) continuously recorded the handle horizontal and vertical positions (100 Hz with 0.013 cm accuracy). The behavioral task was implemented by means of LabVIEW 2018 (National Instruments, Austin, TX). Visual stimuli and handle position feedback (black cross) were projected by a DELL P2219H LCD monitor (60 Hz refresh rate) onto a half-silvered mirror suspended 26 cm above and parallel to the digitizer plane, creating the illusion for the participants that stimuli floated on the plane of the tablet (please see Fig. 1a in Saleri Lunazzi et al., 2023).

### Behavioral task

Participants performed multiple trials of a multi-step decision-making task (Fig. 1). Each trial begins with a black circle (the starting circle, Ø = 3cm) displayed at the bottom of the screen. To initiate a trial, the subject moves the handle in the starting circle and maintains the position for 300ms. A large (Ø = 9cm) circle then appears on the screen (the decision stimulus) for 300ms. It contains 100 green and blue tokens. The ratio between blue and green tokens defines the coherence of the decision stimulus. At this stage, the stimulus informs the subject about the difficulty of the perceptual decision she/he has to make later in the trial, but the dominant color among the tokens is not informative of the correct target to select at the end of the trial. The proportion of tokens of the dominant color is 77% on the first trial of the session.

Then the decision circle disappears and two rectangles are displayed above the starting circle, separated from each other by 10 cm. In each rectangle a text informs the subject about the difficulty of the movement that she/he has to execute in each trial to report the perceptual decision: “1” for an easy movement, executed toward a large visual target (Ø = 3.75cm, with a trial-to-trial variability of 2%), or “5” for a difficult movement, executed toward a small target (Ø = 1.25cm, with a trial-to-trial variability of 2%). The subject has 1s to move the handle in the chosen rectangle and then must hold it for 500ms to validate this choice. She/he then returns to the starting circle and maintains the position for another 500ms to continue the trial.

The decision circle (filled with the 100 tokens) as well as the blue and green movement targets then appear. The movement targets are visual circles displayed 180° apart of the starting circle. Their size depends on the choice of the subject, either large (Ø = 3.75cm) or small (Ø = 1.25cm). The distance between the starting circle center and each movement target center was 10cm, with a trial-to-trial variability of 1.9cm. The dominant color among the tokens (with the same proportion as at the beginning of the trial) now determines the correct target to select.

The subject task is to determine the dominant color in the decision circle, either blue or green. To express this perceptual decision, the participant moves the handle in the lateral target whose color corresponds to her/his choice and maintains this position for 500ms. The dominant color (blue or green) as well as the position of the green and blue movement targets relative to the starting circle are randomized from trial to trial. The maximum decision duration allowed (the time between the decision circle onset and movement onset) is 1s. The maximum movement duration allowed (the time between movement onset and offset) is 750ms.

At the end of the trial, a visual cue informs the subject about the outcome of the trial. The chosen target was surrounded by a green circle if she/he accurately reaches the correct target, and by a red one if she/he accurately reaches the wrong target. The subject earns the number of points corresponding to the chosen difficulty of the movement to execute if the correct target was accurately reached. The goal of the subject is to earn a total of 200 points. If the subject fails to reach or stop in the chosen target (inaccurate movement, whether it is the correct target or not), the number of points chosen at the beginning of the trial is subtracted. Regardless of the movement constraint level chosen by the subject, if she/he selects the wrong target (decision error) with an accurate movement, points are not deducted. To move on to the next trial, the subject moves the handle back in the starting circle and maintains the position for 500ms.

In this experiment, the number of points accumulated by the subject determines the coherence of the decision stimulus. The coherence of the decision stimulus is initially set to 77% at the beginning of the session and it linearly decreases with the accumulation of points, reaching 51% at 200 points. As a consequence, the difficulty of decision progressively increases as the subject gets close to 200 points. We expected to observe with this design a mixture of choices of the difficulty of the movements to be performed to express the perceptual decisions, regardless of the coherence of the decision stimulus, with a bias for the most difficult movements chosen more frequently at the beginning of the session (when perceptual decisions are easier) than at the end.

### Instructions provided to the subjects

To familiarize each participant with the task and with the manipulation of the handle on the tablet, a training phase was proposed prior to the experimental phase per se. During this training phase, subjects performed about 20 training trials where they could choose the difficulty of the movement to make (easy or difficult) and report moderately difficult (63% coherence) perceptual decisions by executing reaching movements to those targets. The training phase was prolonged if subjects required so. During the experimentation phase, each subject was instructed to perform the task described above and they were informed that they needed to earn a total of 200 points to complete the session. Importantly, the 32 subjects who performed the task were not told about the decreasing coherence of the decision stimulus indexed to the accumulation of points. They were also not told about their number of points accumulated after each trial. We informed the subjects that there would be no scheduled breaks during the session, except in case of discomfort or real fatigue. No subject requested a break during their session.

### Data analysis and statistics

Data were collected by means of LabVIEW 2018 (National Instruments, Austin, TX), stored in a database (Microsoft SQL Server 2005, Redmond, WA), and analyzed off-line with custom-written MATLAB scripts (MathWorks, Natick, MA).

Arm movement characteristics were assessed using the subjects’ movement kinematics. Horizontal and vertical arm position data (collected from the handle on the digitizing tablet) were first filtered using a tenth-degree polynomial filter and then differentiated to obtain a velocity profile. Onset and offset of movements were determined using a 3.75 cm/s velocity threshold. Peak velocity and amplitude was determined as the maximum value and the Euclidian distance between movement onset and offset, respectively.

An accurate movement is defined as a movement that reached a target (whether it is the correct target or not) and stayed in it for 500ms. In this report we only refer to movements executed to report the perceptual decisions (not those executed to select the difficulty of the movement to perform to report the perceptual decisions). Decision duration is defined as the time between the onset of the stimulus providing the visual evidence to the subject (the decision circle containing the 100 tokens) to the onset of the movement executed to report the decision. A decision is defined as correct if the correct target is chosen, regardless of the accuracy of the movement.

Chi-squared tests for independence were used to assess the effect of movement difficulty (constrained or less constrained) on individual subjects’ movement and decision accuracy. Wilcoxon rank sum tests were used to assess the effect of movement constraint level on individual subjects’ decision and movement duration. Chi-squared tests for independence were used to test the effect of decision difficulty, evaluated by chronologically grouping trials in 10 quantiles, on individual subjects’ proportion of constrained movement choices. At the population level, Kruskal-Wallis tests were used to test the effect of decision difficulty on the proportion of constrained movement choices. Analyses of covariance (ANCOVAs) were used to assess the effect of decision difficulty, motor constraint level and their interaction on decision accuracy and duration. The significance level of all statistical tests was set at 0.05, and highest levels of significance are reported when appropriate.

## Acknowledgements/Funding

The authors have no financial or non-financial interests to declare. The authors wish to thank Sonia Alouche and Jean-Louis Borach for their administrative assistance, and Frédéric Volland for his expertise during the technical preparation of this experiment. This work is supported by a CNRS/Inserm ATIP/Avenir grant to DT.

## AUTHORS’ CONTRIBUTION

EL, EK and DT designed the experiment EK coded the task

EL collected the data

EL and DT conducted the analyses and prepared the figures

DT wrote the draft of the manuscript

EL, EK and DT revised the draft and approved the final version of the manuscript

## DATA AVAILABILITY STATEMENT

The datasets used and/or analyzed during the current study are available from the corresponding author on reasonable request.

