## Supplemental Figures 1 and 2 for "Humans anticipate the consequences of motor control demands when making perceptual decisions between actions"

### *Supplemental information*

We analyzed the effect of the increasing decision difficulty on subjects' movement accuracy, duration, speed and amplitude, by grouping trials depending on the motor condition chosen by the subjects (small targets, difficult movement vs. large targets, easy movement). We observed a strong effect of the motor condition on the accuracy of participants' movements (ANCOVA, Movement difficulty, F=360, p<0.0001), regardless of the trials performed during the session (Movement difficulty x Trials: F=2.45; p=0.12) (Fig. S1, top-left). We also observed a significant effect of the motor condition chosen by the subject on movement duration (Movement difficulty: F=14, p = 0.0002), as well as a trend for movement durations being influenced by decision difficulty (Trials: F=3; p=0.08), without a significant interaction between the two factors (Movement difficulty x Trials: F=0.7; p=0.4) (Fig. S1, bottom-left). The speed of the movement was not influenced by the motor condition (Movement difficulty, F=0.34, p=0.56) neither with the difficulty of the perceptual decision (Trials, F=3, p=0.06) (Fig. S1, top-right). Finally, we observe that amplitude of movements was larger in the small targets condition compared to the large targets condition (Movement difficulty, F=6.81, p=0.009), and amplitude increased as a function of the evolving decision difficulty (Trials, F=5.9, p=0.02), especially in the small target condition (Movement difficulty x Trials: F=4.3; p=0.04) (Fig. S1, bottom-right).

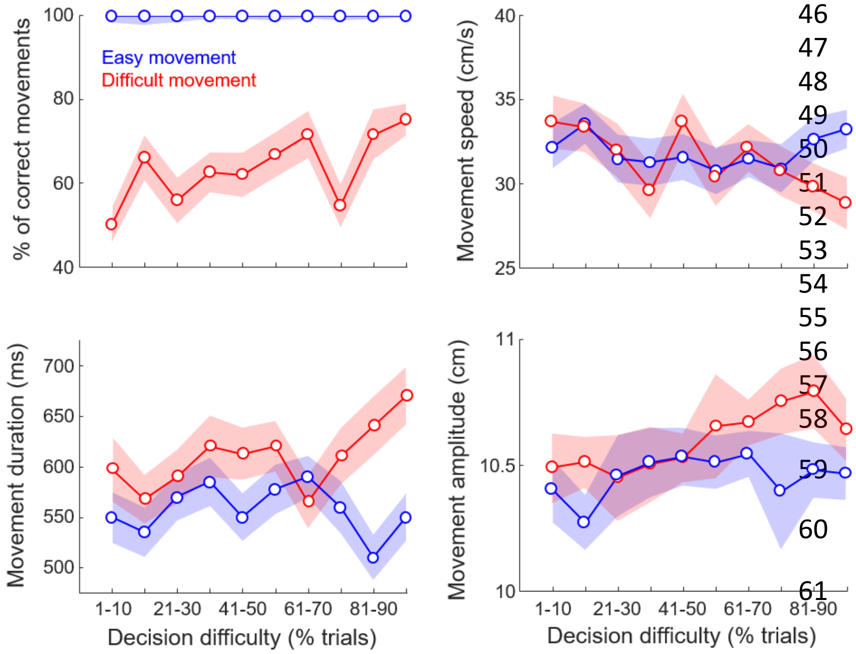


**Figure S1**: Effect of decision difficulty on subjects' movement accuracy (top-left panel), duration (bottom-left), speed (top-right) and amplitude (bottom-right) with trials sorted according to the motor condition chosen by the subjects (red: small targets, difficult movement; blue: large targets, easy movement). Same conventions as in Figure 4C-D in the main manuscript.

Finally, we computed for each trial the duration of what we called the participants' behavior (or the response time), corresponding to the addition of decision and movement durations. We analyzed the effect of the increasing decision difficulty on this metric, by grouping trials depending on the motor condition chosen by the subjects (small targets, difficult movement vs. large targets, easy movement). We observed an effect of the increasing decision difficulty on the duration of participants' behavior (ANCOVA, Trials, F=56.4, p<0.0001), but we found no significant effect of the motor condition (Movement difficulty, F=0.52, p=0.47) as well as of the interaction between the motor condition and the difficulty of the decisions (Trials x Movement difficulty, F=0.43, p=0.5) on behavior duration (Fig. S2).

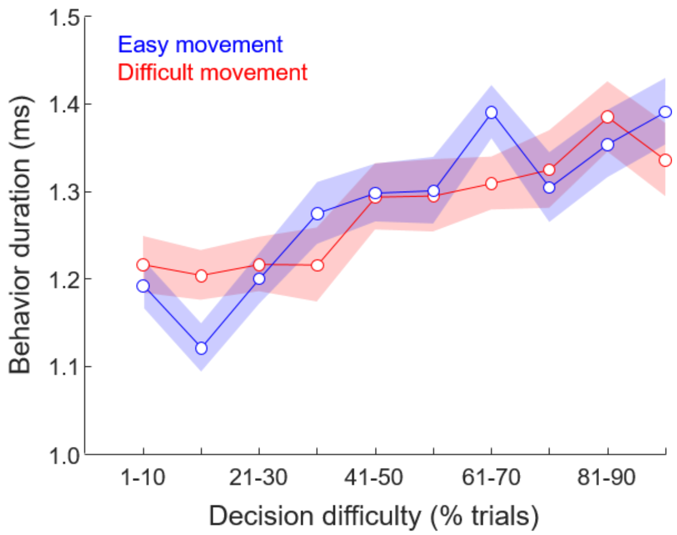


**Figure S2**: Effect of decision difficulty on subjects' behavior (decision and movement) duration, with trials sorted according to the motor condition chosen by the subjects (red: small targets, difficult movement; blue: large targets, easy movement). Same conventions as in Figure S1.
